# Single nucleus transcriptome of a “Super RV” shows increased insulin and angiogenesis signaling

**DOI:** 10.1101/2023.11.28.569092

**Authors:** Diwakar Turaga, Xiao Li, Yi Zhao, Chang-Ru Tsai, Axel Moreira, Edward Hickey, Iki Adachi, James Martin

**Author notes:** These authors contributed equally.

## Abstract

The right ventricle (RV) is one of the four pumping chambers of the heart, pumping blood to the lungs. In severe forms of congenital heart disease and pulmonary hypertension, the RV is made to pump into the systemic circulation. Such systemic RVs typically display early failure due to pressure overload. In rare cases a systemic RV persists into later decades of life – colloquially called a ‘Super RV’. Here we present the single-nucleus transcriptome of a systemic RV from a 60-year-old with congenitally corrected transposition of great arteries (ccTGA). Our data shows two specific signaling pathways enriched in the ccTGA RV myocardium. First, we show increased insulin like growth factor (IGF1) signaling within the systemic RV myocardium: there is increased expression of the main receptor IGFR1 within the cardiomyocytes, and IGF1 ligands within the cardiofibroblasts and macrophages. Second, we find increased VEGF and Wnt9 ligand expression in cardiomyocytes and increased VEGF1R and Wnt9 receptors in endothelial cells, which are implicated in angiogenesis. We show that increased insulin and angiogenesis signaling are potentially beneficial RV adaptations to increased pressure overload. This study of an adult systemic RV provides an important framework for understanding RV remodeling to systemic pressures in congenital heart disease and pulmonary hypertension.

## INTRODUCTION

In the normal heart the right ventricle (RV) is primed to pump blood to the lungs and lives in a lower pressure environment. The RV is made to pump into systemic or higher pressures in specific forms of congenital heart disease (CHD) and severe cases of pulmonary hypertension (PH). The systemic RV must undergo adaptive mechanisms to pump against the pressure overload. In CHD such systemic RVs encompasses 10-12% of all forms CHD [1], and include congenitally corrected TGA (ccTGA), hypoplastic left heart syndrome and d-transposition of great arteries who underwent atrial switch. In pulmonary hypertension there is remodeling of pulmonary vasculature and increase in RV afterload, with severe cases of pulmonary hypertension seeing systemic or supra-systemic RV pressures [2]. These systemic RVs fail early, and the reason is thought to be multi-factorial – including distinct RV tissue architecture, shape and function, tricuspid valve abnormalities or inability to adapt to pressure overload.

ccTGA is a rare structural heart disease with suggested incidence of 1 in 33,000 live births [3]. It is a unique congenital heart lesion where despite atrioventricular and ventriculoarterial discordance, there is a physiologically normal circulation [4]. The natural history of unrepaired ccTGA without significant intracardiac lesions is progressive failure of systemic RV [5], though the timing of presentation is variable and many patients have functional lives [6]. The systemic RV leads to manifestation of ventricular dysfunction and symptoms during the 4^th^ or 5^th^ decade of life [7]. The majority of ccTGA patients have associated lesions such as ventricular septal defects, pulmonary stenosis, ebsteinoid tricuspid valve and abnormal conduction system. Early diagnosis of ccTGA allows for the double switch procedure and accompanying lesions to be addressed surgically. But a late presentation of ccTGA is usually in the setting of clinical heart failure, and such end stage heart failure often necessitates placement of a ventricular assist device (VAD) and/or heart transplantation.

We present a rare 60-year-old patient with ccTGA, without any accompanying intracardiac lesions, who presented with systemic RV failure and underwent placement of a VAD. We performed single nucleus RNA sequencing (snRNA-seq) to identify gene expression changes in this systemic RV compared to controls [Fig 1A].

**Figure 1:**
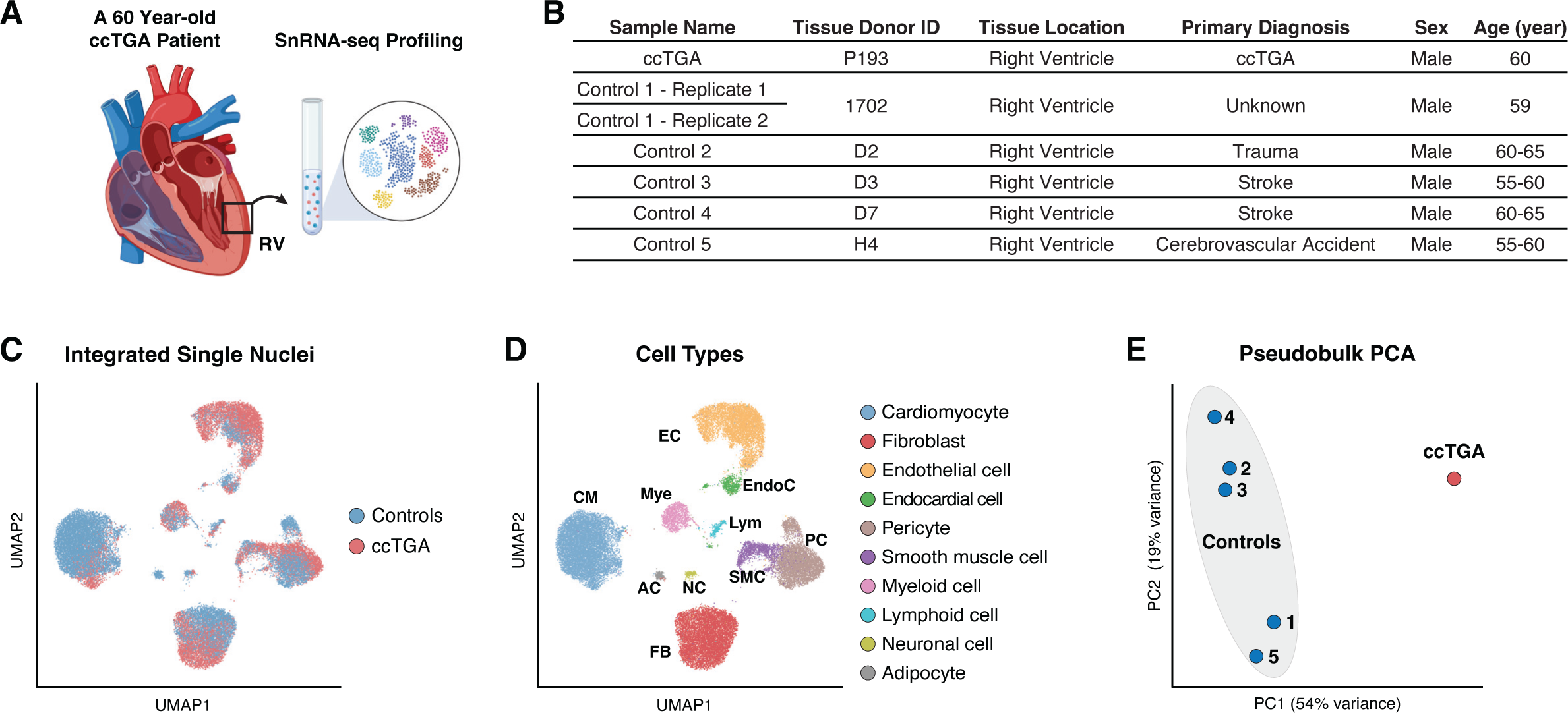
Single nucleus profiling of ccTGA RV. **[A]** Schematic illustrating design of the snRNA-seq experiment. Right ventricular biopsy was collected from the systemic right ventricle of a 60-year-old patient with ccTGA at the time of VAD placement. **[B]** Table of clinical data of index patient and controls. **[C]** Global uniform manifold approximation and projection (UMAP) of the integrated dataset nuclei from ccTGA RV combined with nuclei of RV from controls. **[D]** Global UMAP showing 10 distinct cardiac cell types based on unbiased clustering and reference-assisted annotation. **[E]** Principal component analysis (PCA) bubble plot of pseudobulk snRNA-seq colored by index patient (red) and control patients (blue).

## RESULTS

This study evaluates a 60-year-old patient with ccTGA who underwent placement of a VAD due to worsening systemic RV failure. We profiled the RV myocardium from this patient using snRNA-seq and compared the gene expression landscape with publicly available snRNA-seq data from RV myocardium of five age-matched donor RV tissues as controls [Fig 1B] [8, 9].

We integrated the single-nuclei transcriptome from ccTGA patient with the five controls. We utilized a bioinformatic pipeline that incorporates rigorous quality controls, inter-batch integration, and reference-assisted annotation [Fig S1A, S1B]. We identified 10 cell clusters after filtering doublets and ambient RNA-contaminated nuclei [Fig 1C, S1C, Table S2]. Assisted by established adult heart atlas data, we annotated each cluster based on their transcriptional signatures [Fig S1D]. All major cardiac cell types were recovered, including cardiomyocytes (CMs), cardiac fibroblasts (FBs), endothelial cells (ECs), vascular smooth muscle cells (SMCs), and a myeloid population which primarily consists of macrophages (MPs) [Fig 1D]. Principal component analysis on pseudo-bulk transcriptome revealed clear global transcriptional distinction between ccTGA patient and controls [Fig 1E].

Unbiased graph-based iterative clustering on CMs from the complete cardiac tissue snRNA-seq dataset uncovered five CM clusters [Fig 2A, Table S3]. CM1, CM3 and CM4 contained nuclei from both ccTGA and control patients. CM2 predominantly contained nuclei from control patients and CM5 was significantly enriched in the ccTGA patient [Fig 2B, C].

**Figure 2:**
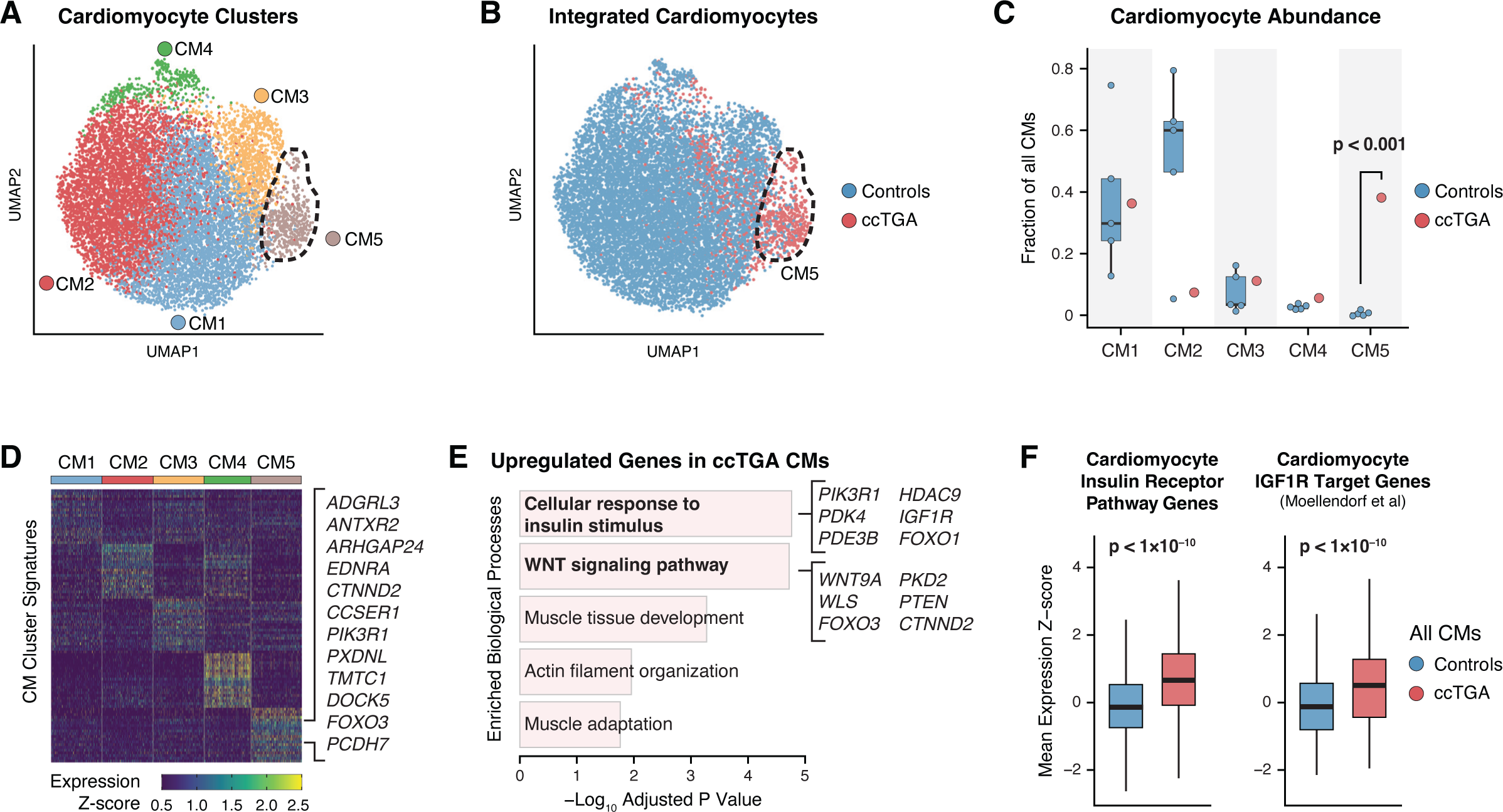
Increased insulin signaling in ccTGA specific CMs. **[A]** UMAP embedding of clustering of CMs revealing five distinct clusters. **[B]** UMAP embedding of CMs with ccTGA nuclei in red and controls in blue. **[C]** Fractional abundance ccTGA and control nuclei for each CM cluster. **[D]** Heatmap of average expression of genes for each CM cluster, with top genes of CM5 highlighted. **[E]** Gene Ontology analysis of ccTGA CM upregulated genes, including genes in insulin signaling and WNT signaling pathways. **[F]** Left, Gene expression of human CM insulin receptor pathway genes for controls compared to ccTGA. Right, Gene expression of mouse CM Igf1r target genes for controls compared to ccTGA.

We focused our analysis on CM5, a cardiomyocyte cell state highly associated with the ccTGA patient. The genes differentially expressed in this cluster include IGF1R, PIK3R1, and FOXO3 [Fig 2D, Table S3]. Gene Ontology (GO) analysis of the CM5 signature genes was significant for upregulation of genes involved in cellular response to insulin stimulation and Wnt signaling pathway [Fig 2E, Table S4]. We found significantly increased expression of human CM insulin receptor pathways genes in ccTGA CMs when compared to those in controls [Fig 2F]. IGF1R-activated downstream genes identified in a CM-specific Igf1r deletion mouse model [10] were also significantly upregulated in ccTGA CMs [Fig 2F], further demonstrating the hyperactivity of IGF1 signaling in ccTGA CMs. In addition, we found increased non-canonical WNT signaling pathway enriched in ccTGA CMs, including ligand WNT9A and transporter WLS [Fig 2E, Table S4].

CellChat, a cell-cell communication inference suite [11], was used to model the significantly enriched ligands sent by non-CM cell types and received by CMs in the ccTGA RV [Fig 3A]. One particularly strong interaction is IGF1 from fibroblasts and macrophages interacting with IGF1R (the main receptor for IGF1) on the CMs [Fig 3A, B].

**Figure 3:**
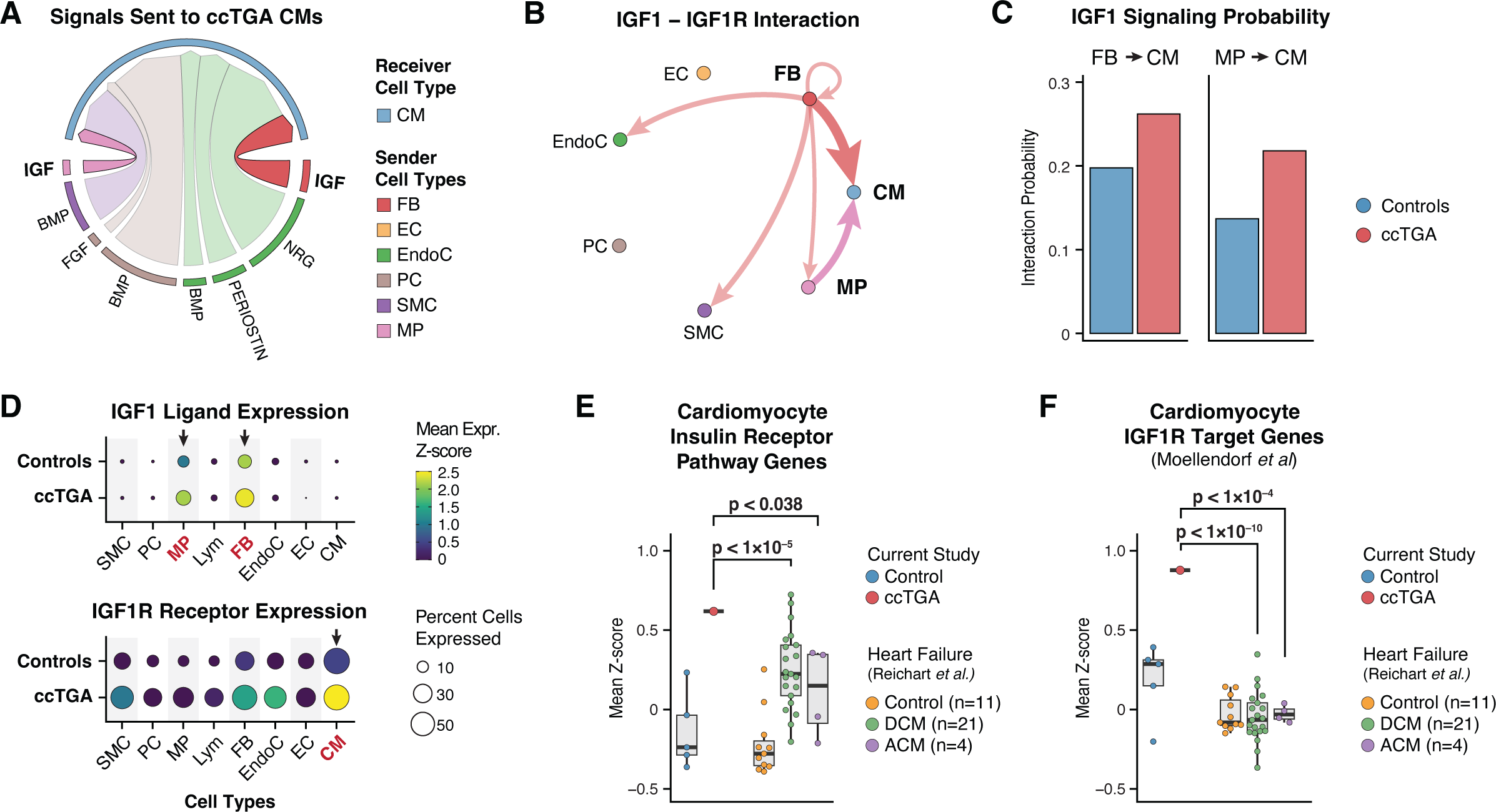
IGF1-IGF1R signaling from cardiac fibroblasts / tissue resident macrophages to cardiomyocytes. **[A]** Chord diagrams of predicted ligand-receptor interactions between ccTGA CMs and other cell types (FB, cardiac fibroblasts; EC, endothelial cells; EndoC, endocardial cells; PC, pericytes; SMC, smooth muscle cells; Mye, myeloid cells). **[B]** Significant IGF1-IGF1R signaling predicted between ccTGA CMs and other cell types. Vertex diagram illustrates the direction (arrow) and strength (thickness) of interactions. **[C]** IGF1 signaling between FBs and CMs (left) and macrophages and CMs (right) for ccTGA and control CMs. **[D]** IGF1 ligand expression (top) and IGF1R Receptor expression within different cell types. **[E]** Gene expression of human CM insulin receptor pathway genes for ccTGA compared to dilated cardiomyopathy (DCM) and arrhythmogenic cardiomyopathy (ACM). **[F]** IGF1R target genes for ccTGA CMs compared to DCM and ACM myocardium.

The FB to CM and MP to CM signaling probabilities were both increased in ccTGA patient [Fig 3C]. We see increased IGF1 ligand expression in myeloid and fibroblasts in ccTGA patient as well as complementary increased IGF1R receptor expression in ccTGA CMs [Fig 3D]. We then asked the question whether failing RV leads to increased IGF1 signaling. We compared ccTGA myocardium with published dilated cardiomyopathy (DCM) and arrthymogenic cardiomyopathy (ACM) single-nucleus data from Reichart et al. [12]. We show that even though human cardiomyocyte insulin pathway receptor pathway genes are increased in DCM compared to controls, ccTGA CMs have a higher increase than DCM CMs [Fig 3E]. When we compare Igf1r receptor target genes (based on mouse study [10]) we see increased IGF1 signaling in ccTGA patient compared to DCM and ACM patients [Fig 3F].

To further investigate the sources of IGF1 ligands, we clustered FBs and myeloid cells at a finer resolution. Our clustering of FBs uncovered four clusters [Fig 4A, Table S5]. FB4 population was significantly enriched with nuclei from ccTGA patient [Fig 4B, C]. This cluster expressed high levels of genes associated with activated cardiac fibroblasts, such as POSTN and CCN2 [Fig 4D, Table S5]. This was also the cluster with highest IGF1 expression [Fig 4D].

**Figure 4:**
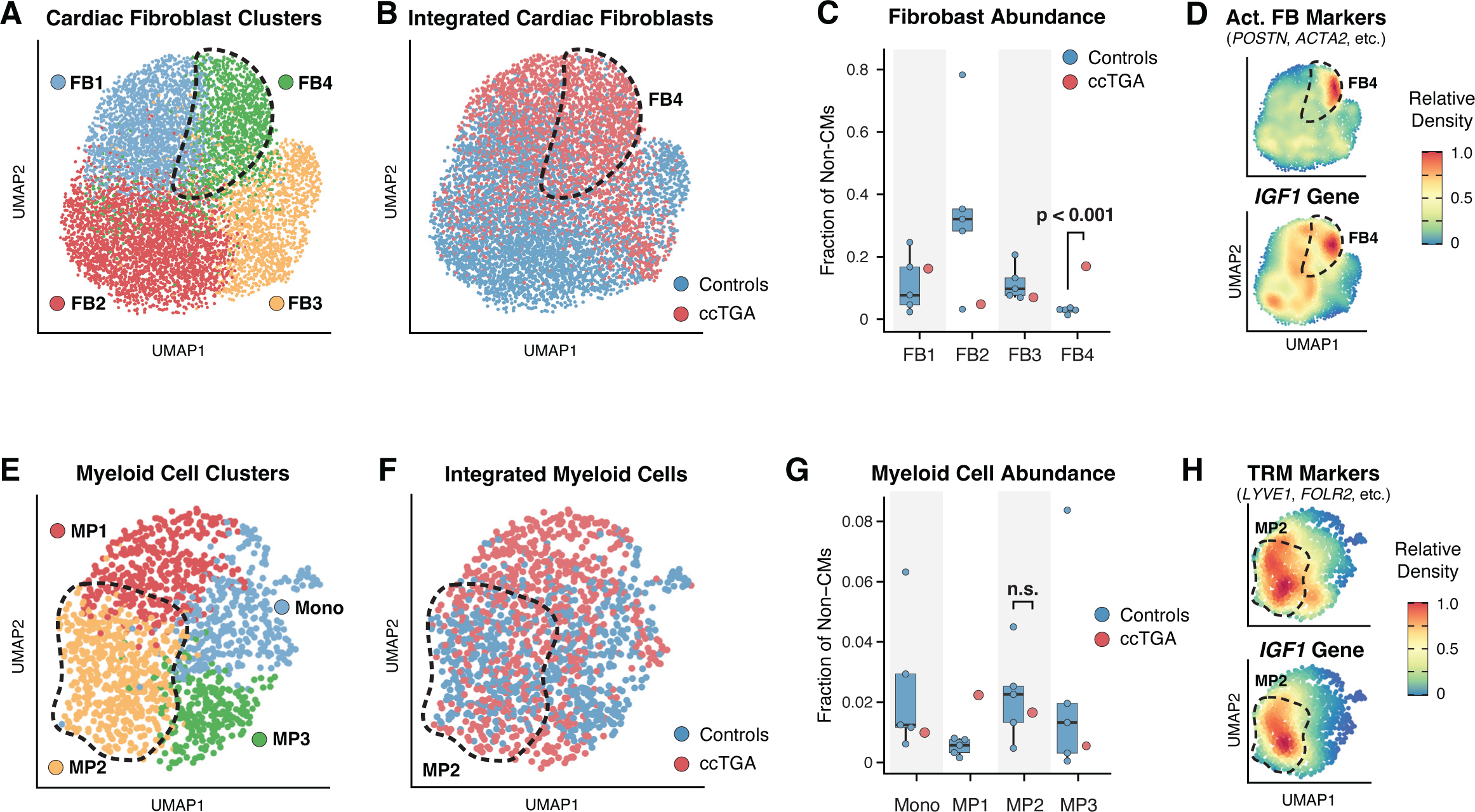
Cardiofibroblasts and tissue resident macrophages in ccTGA myocardium show increased IGF1 gene expression. **[A]** UMAP embedding of reiterative clustering of FBs revealing four distinct clusters. **[B]** UMAP embedding of FBs with ccTGA nuclei in red and controls in blue. **[C]** Fractional abundance ccTGA and control nuclei for each FB cluster. **[D]** Relative density of activated fibroblast markers (Act. FB Markers) and IGF1 gene. **[E]** UMAP embedding of clustering of myeloid cells reveals four distinct clusters. **[F]** UMAP embedding of myeloid cells with ccTGA nuclei in red and controls in blue. **[G]** Fractional abundance ccTGA and control nuclei for each myeloid cell cluster. **[H]** Top, Relative density of tissue resident macrophage (TRM) markers. Bottom, Relative density of IGF1 gene expression.

Our clustering of myeloid cells uncovered four main clusters, each composed of ccTGA and control nuclei [Fig 4E, 4F, Table S6]. We did not observe significant compositional change of MP2 population when comparing ccTGA with controls [Fig 4G]. The MP2 cluster strongly expressed tissue-resident macrophage (TRM) signature genes [Fig 4H], such as LYVE1 and MRC1. MP2 cells also specifically expressed high level of IGF1 [Fig 4H].

Next, we sought to examine the cell-to-cell communications initiated by CMs and received by non-CMs in the ccTGA RV. Using CellChat, we identified EC as a major cell type that receives signals from CMs in the ccTGA RV [Fig 5A]. Specifically, we identified VEGF signaling as the most significant interactions [Fig 5A]. WNT signaling was also predicted as a significant interaction sent from CM to ECs, which is aligned with the ccTGA-enriched CM5 signature [Fig 5A, 2E].

**Figure 5:**
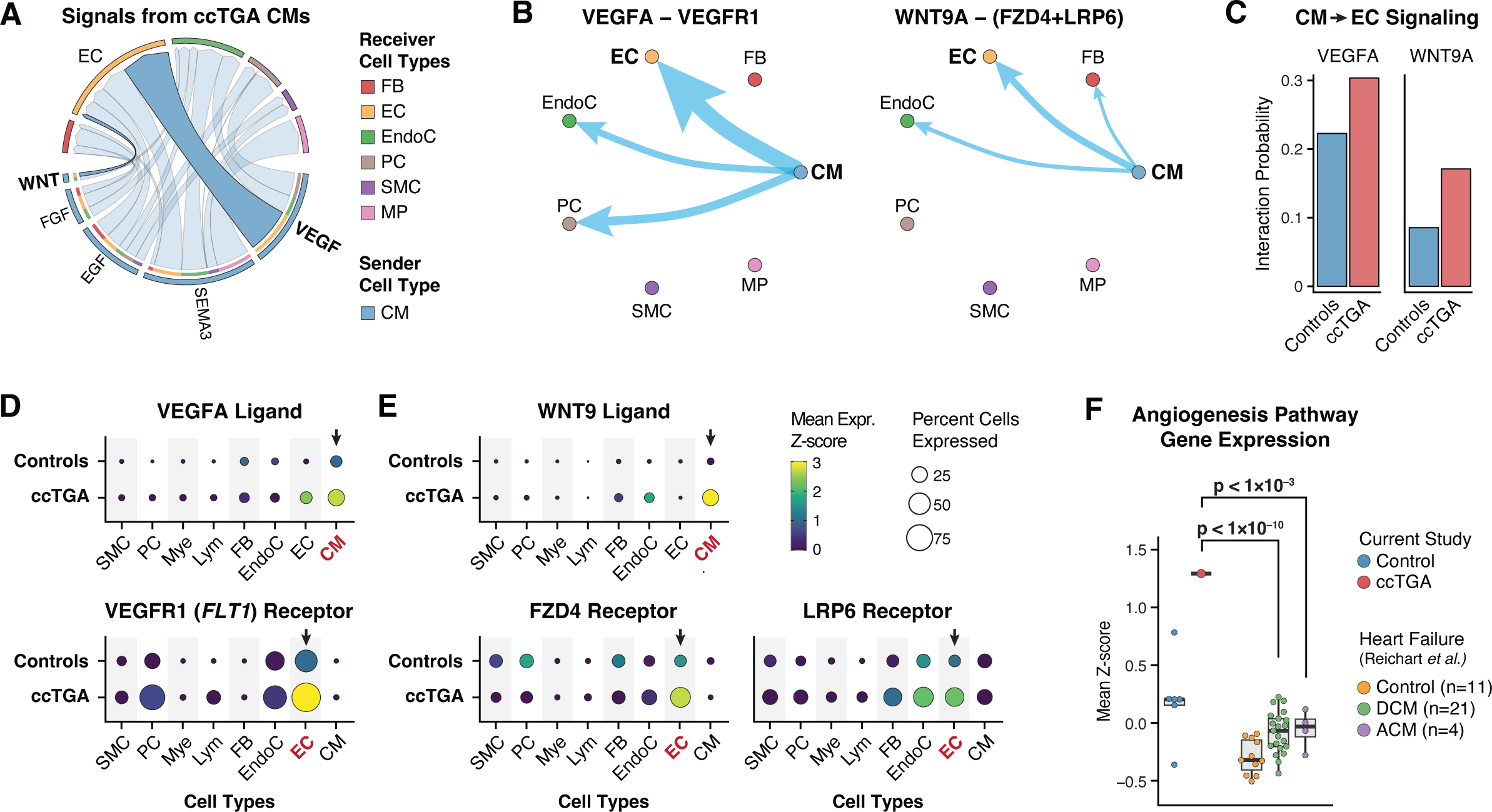
VEGF and WNT signaling from ccTGA CMs to Endothelial Cells (ECs). **[A]** Chord diagrams of predicted ligand-receptor interactions from ccTGA CMs to other cell types (FB, cardiac fibroblasts; EC, endothelial cells; EndoC, endocardial cells; PC, pericytes; SMC, smooth muscle cells; MP, macrophages). **[B]** Significant VEGFA-VEGFR1 and WNT9-[FZD4+LRP6] signaling predicted from CMs to ECs. Vertex diagram illustrates the direction (arrow) and strength (thickness) of interactions. **[C]** CM to EC signaling strength for VEGFA and WNT9A signals in ccTGA vs controls. **[D]** VEGFA ligand (top) and VEGFR1 receptor (bottom) expression within different cell types. **[E]** WNT9 ligand (top) and FZD4 receptor (bottom) expression within different cell types. **[F]** Angiogenesis pathway gene expression for genes from ccTGA compared to controls, DCM and ACM myocardium.

Detailed examination revealed that VEGF signaling was mediated by VEGFA-VEGFR1 interactions, and the WNT signaling by WNT9A-FZD4/LRP6 interactions [Fig 5B]. Both interactions were most strongly enriched between CMs and ECs [Fig 5B]. The probabilities of these communications were both increased in ccTGA patient when compared with the controls [Fig 5C].

The expression of ligand VEGFA is highest in ccTGA CMs over other cell types, and that of receptor VEGFR1 is highest in ccTGA ECs, further corroborating the CellChat results. [Fig 5D]. Similarly, the expression of WNT9A ligand with receptors FZD4 and LRP6 are also highly specific between CMs and ECs in ccTGA compared to controls [Fig 5E].

VEGFA is a major driver of myocardial angiogenesis. WNT9A has also been implicated in promoting angiogenesis in other systems. We reason that the enriched interactions of these signals in the ccTGA heart may promote angiogenesis, which has been considered as an adaptive remolding feature during HF progression. To test this, we compared ccTGA myocardium with published DCM and ACM single-nucleus data [12]. We show that angiogenesis pathway genes collectively upregulated significantly in ccTGA ECs compared to those in control hearts, as well as in failing DCM and ACM hearts [Fig 5F]. This result suggests that angiogenesis activity is likely a unique feature of this ccTGA RV, rather than a general phenotype associated with myocardial remodeling in failing hearts.

To further investigate the increased angiogenesis phenotype, we performed detailed sub-clustering of ECs, which revealed 5 main EC clusters (EC1 through EC5) [Fig 6A]. Their distinct gene expression signatures suggest that they may correspond to specific vessel types, such as capillary (EC1), artery (EC2), vein (EC3), lymphatic (EC4) and endocardial (EC5) [Fig 6A, Table S7]. Major blood vessel cells EC1-3 are all significantly enriched in ccTGA, while EC2 displayed the strongest increase [Fig 6B, C]. In the integrated data including both ccTGA and controls, EC2 population highly expresses receptors VEGFR1 (FLT1 gene), FZD4 and LRP6, suggesting that it is the main cell population that receives the angiogenesis signaling from CMs. Consistently, EC2 cells also showed highest mean expression of angiogenesis pathway genes [Fig 6D].

**Figure 6:**
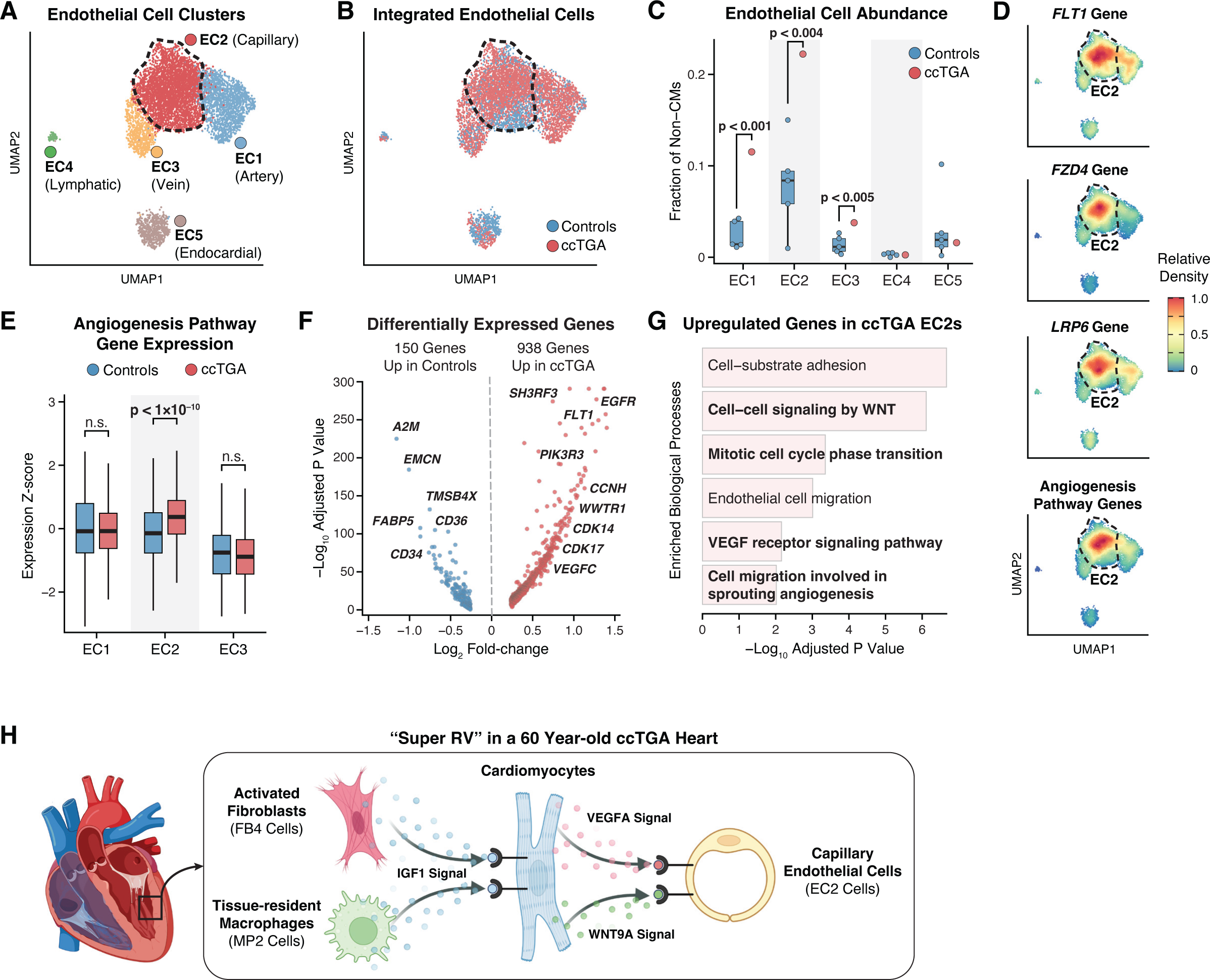
Increased angiogenesis gene expression in ccTGA ECs. **[A]** UMAP embedding of clustering of ECs revealing five distinct clusters, each corresponding to artery, capillary, vein, lymphatic and endocardial vessels. **[B]** UMAP embedding of ECs with ccTGA nuclei in red and controls in blue. **[C]** Fractional abundance ccTGA and control nuclei for each EC cluster. **[D]** Relative density in EC2 for VEGF1 receptor (FLT1), FXD4, LRP6, and mean expression of angiogenesis genes. **[E]** Angiogenesis pathway gene expression for EC1, EC2 and EC3 for controls vs. ccTGA. **[F]** Volcano plot depicting differentially expressed genes for EC2, with 938 genes up in ccTGA and 150 genes up in controls. **[G]** Gene Ontology analysis of ccTGA EC2s upregulated genes. **[H]** Model of ccTGA RV remodeling showing two enriched pathways: (i) activated fibroblasts and tissue-resident macrophages have increased expression of IGF1 ligands which are detected by cardiomyocytes expressing increased IGF1R receptor, and (ii) pro-angiogenesis signaling between CM and ECs via VEGF and Wnt9 pathways.

When compared to the controls, EC2 cells from ccTGA RV showed significantly upregulated expression of angiogenesis pathway genes [Fig 6E]. Differential gene expression analysis of EC2 population identified 150 genes that are downregulated and 938 upregulated in ccTGA compared to controls [Fig 6F]. The upregulated genes are enriched in biological processes such as cell−cell signaling by WNT, mitotic cell cycle, VEGF receptor signaling, and importantly, cell migration in sprouting angiogenesis [Fig 6G]. These functional inferences all point to the involvement of CM-EC signaling axis in myocardial angiogenesis.

Overall, our ccTGA data highlights two adaptive mechanisms exhibited by ccTGA RV to systemic pressures. First, we find upregulation of IGF1 signaling originating from FB4 (the activated cardiac fibroblasts) and MP2 (the tissue-resident macrophages), and the complementary increased IGF1R receptors in CMs of ccTGA myocardium [Fig 6H]. Second, we find upregulation of VEGF and Wnt9 in CMs, and complementary increase in VEGFR1, FZD4, LRP6 in ECs, which promote angiogenesis [Fig 6H].

## DISCUSSION

In this report we present the first single-nucleus transcriptome of an adult systemic RV. Congenital heart disease lesions such as HLHS, ccTGA, d-TGA with atrial switch and severe forms of pulmonary hypertension have systemic RVs. These RVs fail due to increased pressure load on the RV, but some rare RVs tolerate the increased loading conditions. These RVs are colloquially called ‘super RVs’. There is strong interest in the research community to understand why some RVs succeed (super RVs) while others fail. In this study we perform an in-depth gene expression analysis of one such ‘super RV’. Understanding the specific remodeling adaptations of the ‘super RV’ will pave the way for new therapeutic targets and enhance the longevity of failing systemic RVs.

In this paper we show two specific molecular adaptations in the super RV. First, we show that ccTGA CMs have a distinct upregulation of IGF1 receptor. Complementarily, ccTGA FBs and MP2s show an upregulation of IGF1 ligands. IGF1 signaling regulates contractility, metabolism, hypertrophy in the heart [13]. IGF1 signaling modulates the cardiac responses to physiological and pathological stressors. Altered IGF1 signaling in the heart may contribute to the pathophysiology of ventricular remodeling and heart failure progression. Resident macrophage IGF1 has recently been shown to play an important role in adaptive cardiomyocyte growth [14]. IGF1 signaling upregulation is seen in the RVs of published human DCM data set. On direct comparison of ccTGA with published DCM data set we find that IGF1 upregulation is higher in ccTGA than in DCM data sets, suggesting that IGF1 upregulation is not entirely due to heart failure phenotype of ccTGA. Therefore, we propose that IGF1 signaling upregulation leads to favorable remodeling of a systemic RV.

The second molecular adaptation we identify is increased in angiogenesis signaling. We show increased VEGF and Wnt9 ligand expression in ccTGA CMs and corresponding increase in VEGF and Wnt9 receptors in ECs. VEGF and Wnt signaling have been shown to be intricately involved in angiogenesis [15]. Angiogenesis is a known adaptation of myocardium to increased stress. Therefore, increased VEGF and Wnt9 signaling may be an adaptative mechanism in the systemic RV.

This data set is an important contribution to our understanding of RV remodeling in adult heart failure, CHD and pulmonary hypertension. Future snRNA-seq studies of pediatric and adult myocardium will be able to use this data set as a comparison in to increase our understanding of favorable and adverse remodeling of RVs.

A major limitation of this study is that it includes RV from only one patient with ccTGA. ccTGA is a rare disease and finding an adult with an isolated ccTGA, is even rarer. We were very fortunate to obtain viable RV myocardium from one such a patient. We compared our tissues against published control RV tissues. In the future, a multi-center study will be needed to obtain myocardial tissues from adults with systemic RVs.

## METHODS

### Research ethics for donated tissue

Cardiac tissue sample used in this study was collected during cardiothoracic surgery performed at Texas Children’s Hospital (Houston, Texas). The protocols for the procurement and use of this patient sample were approved by the Institutional Review Board for Baylor College of Medicine and Affiliated Hospitals (Protocol Number H-26502). With the help of the Heart Center Biorepository at Texas Children’s Hospital, consent was obtained from the patient.

### ccTGA patient information

Patient is a 60-year-old male with congenitally corrected TGA complicated by complete heart block. He had undergone placement of dual chamber transvenous pacemaker. He also underwent multiple cardioversions for atrial fibrillation. He developed systemic RV systolic failure and underwent placement of destination VAD.

### Sample Collection and Nuclear Isolation

Cardiac tissue was collected in the operating room during VAD placement. The anatomic location of tissue collected is RV free wall. Cardiac tissue sample was kept in cold saline on ice during transfer to the laboratory for preservation. Cardiac tissue sample was carefully dissected into multiple aliquots, which were flash-frozen and stored at –80 °C Nuclear isolation was performed as described previously [16, 17]. Briefly, frozen cardiac tissue was dissociated by using a Dounce homogenizer. Single nuclei were isolated via fluorescence-activated cell sorting (FACS).

### Single-nucleus RNA Sequencing

SnRNA-seq was performed by using the 10X Genomics platform. Isolated nuclei were loaded into the 10X Genomics Chromium Controller to obtain the gel beads in emulsion. The sequencing libraries were then prepared according to the manufacturer’s protocols for the Single-cell 3’ Reagents Kits v3. Sequencing was performed by using the NovaSeq 6000 system.

## CONTROLS

Controls tissue data were obtained from the following published data sets. [8, 9]. Data generated in RVs from males with 55-65 year of age were selected as controls. Control data were downloaded in the format of raw sequencing reads and were re-processed with the ccTGA data using the same pipeline.

### SnRNA-seq data processing and integration

All newly generated and published snRNA-seq data sets were processed using a uniformed pipeline. Raw sequencing reads were aligned to the genome (build GRCh38) using the 10X Genomics toolkit CellRanger version 5.0.1 (cellranger count) with --include-introns set to true. All other parameters were left as defaults. Quality control metrics generated by CellRanger were inspected for each library. To remove background signals from ambient transcripts, the raw UMI count matrices were further processed by CellBender version 0.1.0 (cellbender remove-background) with --total-droplets-included = 25000, --low-count-threshold = 15, and --epochs = 200. To minimize the loss of valid cell barcodes called by CellRanger, we also set --expected-cells at 1.5 times of CellRanger output nuclei number. The output matrices from CellBender were filtered to only include valid cell barcodes that were also identified by CellRanger. Additional quality controls at single nucleus level were performed for each library. Briefly, we first identified low-quality nuclei based on fixed cut-offs of UMI count per nucleus > 200, gene count per nucleus > 150 and mitochondria gene-derived UMI < 5%. Then, a set of dynamic cut-offs based on per-library data distribution were calculated, which is essential to account for heterogeneity between samples. In brief, for each library, an upper and lower bound were set at the 75th percentile plus 1.5 times the interquartile range (IQR) and the 25th percentile minus 1.5 times IQR, respectively, for UMI count and gene count per nuclei. Next, the remaining nuclei were evaluated by the Scrublet tool [18] to identify potential doublets, with parameters expected_doublet_rate = 0.1 and call_doublets threshold = 0.25. Finally, we integrated all samples and corrected batch-effect using a deep generative models scANVI [19]. The scANVI latent space was reduced to generate the final global UMAP embedding and subsequent subcluster UMAP embeddings for CMs, FBs and myeloid cells.

### Clustering and annotation

We applied FindNeighbors function of the Seurat version 4.2.0 package to generate the shared nearest-neighbor graph (SNN) using the scANVI latent space. We defined clusters based on the SNN using Louvain algorithm with an optimized resolution of 0.5. As the scANVI model was trained with the Litviňuková, M. et al data set as a reference [9], we examined the predict cell type identities for each cell cluster. Based on both scANVI predicted labels and the expression of known cardiac cell type marker genes, we labeled all main clusters. Subsequent reclustering of each major cell type was performed using an iterative approach. Within each major cell type, subclusters enriched with previously called doublets were examined for the expression of main cell type marker genes and were collectively labeled as ambiguous cells and were excluded from all downstream analysis. The optimal resolutions were determined by clear separation in the UMAP dimension, as well as robust identification of >30 significantly differentially expressed genes across subclusters.

### Principal component analysis

For global comparison of controls and ccTGA cells, principal components were calculated at the pseudobulk level, using the top 1000 highly variable genes. Pseudobulk expression matrix was calculated by summing UMI for each gene across all cells in each sample, using AverageExpression function in Seurat. Then the matrix was normalized, and the principal components were calculated with DEseq2 (version 1.32.0) [20].

### Cell composition analysis

The composition of CMs and FBs was calculated for each of the 5 control and ccTGA samples. For each sample, the cell count of each CM and FB cluster was normalized against the total CM and FB cell count, respectively. Statistical significance was assessed using a two-tailed one-sample t-test to determine if the cell abundances in ccTGA were statistically significant compared to that in controls.

### Gene signature scoring

The Seurat ‘‘AddModuleScore’’ function was used to calculate mean expression Z-scores of various sets of gene signatures. This method scores the expression of a given set of genes by normalizing the expression levels against a set of randomly selected background genes with similar expression levels. The size of the background gene pool is typically 10 times that of the input gene set. Signature scores that fit gaussian distribution across cells were transformed into Z-scores for plotting and statistical significance testing. Specifically, the scores for the “Human Cardiomyocyte Insulin Receptor Pathway Genes” were calculated based on a curated gene set consisting of 51 genes from MSigDB gene set #M7955 (refer to Fig 1K, left). To determine the scores for the “Mouse Cardiomyocyte IGF1R Target Genes,” we reanalyzed microarray data from Moellendorf et al [10]. Using GEO2R, we identified 22 significantly differentially expressed genes that are IGF1R target genes, namely, SULT4A1, CNOT4, ADAMTS16, SOWAHB, KCNU1, LRRC28, PSMA8, PSPN, RBMS3, RNASE13, FAM227B, REM2, HPCAL4, PTGER2, WSCD1, IGSF5, PRDM6, SPATA9, ACACA, FBXL21P, SCARF1, ACSM3. These genes were found to be significantly downregulated in MerCreMer;Igf1r^LoxP/LoxP^ mice compared to MerCreMer control mice. We subsequently used the human orthologs of this gene set to calculate their mean expression in ccTGA and control CMs (refer to Fig 1K, right). For the “Activated FB” score (refer to Fig 2H, top), we utilized established activated cardiac fibroblast signature genes, namely POSTN, ACTA2, DDR2, PDGFA, and COL1A1 (10.1038/nrcardio.2017.57). To calculate the “Tissue-resident Macrophage” score (refer to Fig 2J, top), we employed the genes LYVE1, TIMD4, and FOLR2, which have been previously reported as canonical markers for cardiac tissue-resident macrophages [21]. Statistical significance of expression Z-scores were determined by two-tailed t-test.

### GO and pathway enrichment analysis

GO and pathway were performed using ClusterProfilers [22]. We extracted enriched GO terms in the biological process category. All enriched terms were filtered at a threshold of false discovery rate (FDR, Benjamini-Hoch bergcorrection) at < 0.05.

### Cell-cell communication

A systematic analysis of cell communication was based on the network analysis and pattern recognition approaches provided by CellChat (version 1.5.0) [11]. The standard workflow was used to predict major signaling inputs and outputs of cells and how these cells and signals coordinate with each other for functions. Subsequently, signaling pathways were classified, and conserved and context-specific pathways were depicted between control and ccTGA cells.

## ACKNOWLEDGMENTS

Authors would like to thank Lalita Wadhwa (Texas Children Hospital Heart Center Biorepository) and Alon Azares (FACS core of Texas Heart Institute).

## AUTHOR CONTRIBUTION

Conceptualization: DT, XL, JFM. Human sample and data acquisition: DT, CR-T, AM, EJH, IA. Data analysis and investigation: DT, XL, YZ. Writing manuscript: DT, XL, JFM.

---

**Supplemental Figure 1:**
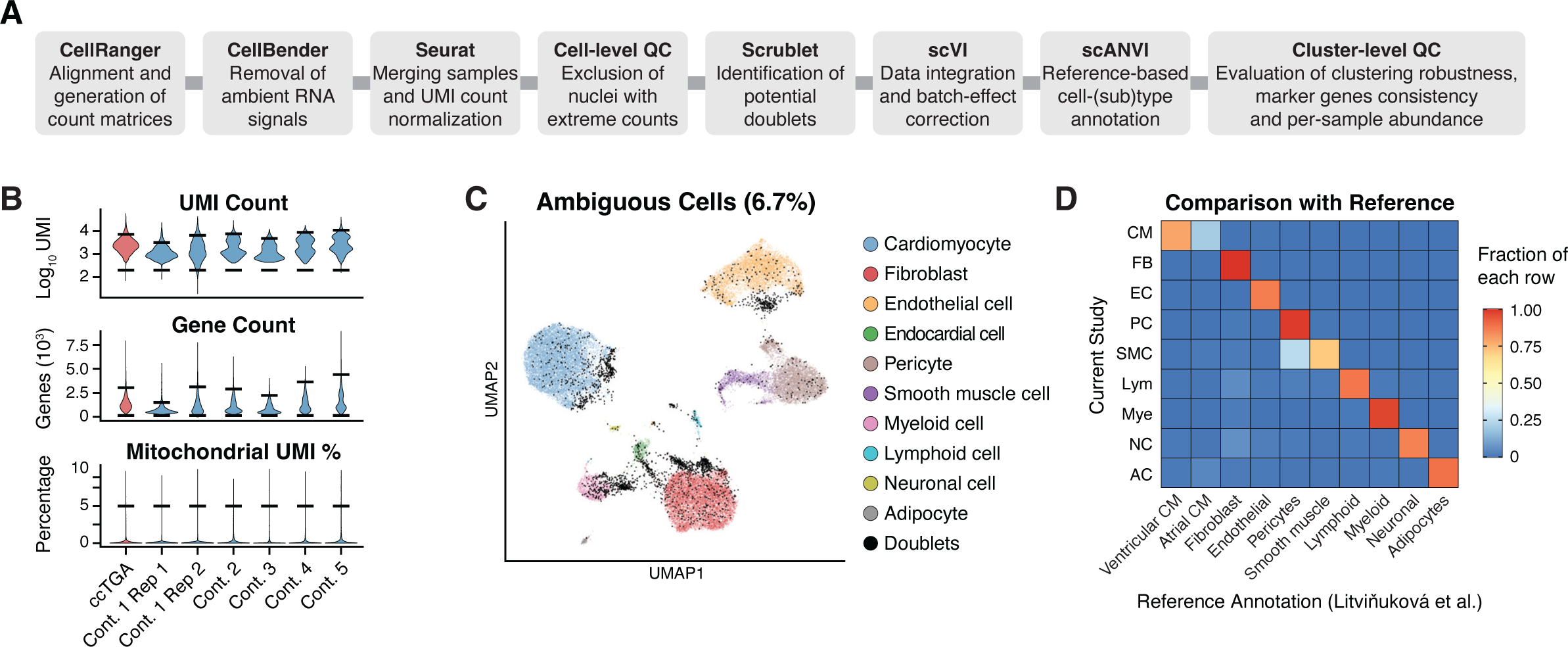
**[A]** A schematic illustrating the key steps of the computational pipeline used for snRNA-seq data processing. [**B**] Violin plots of key quality control metrics of each snRNA-seq library. Black bars indicate the library-specific thresholds. [**C**] A pre-integration UMAP highlighting the low-quality nuclei filtered out by quality controls. [**D**] Heat map of mean expression of reference cell type markers in each of the 9 cell types classified in the integrated data set. Color indicates fraction of cells express the marker genes.

